# Large variations in afforestation-related climate cooling and warming effects across short distances

**DOI:** 10.1101/2022.09.18.508428

**Authors:** Shani Rohatyn, Eyal Rotenberg, Fyodor Tatarinov, Yohay Carmel, Dan Yakir

## Abstract

Climate-related benefits of afforestation depend on the balance of the often-contrasting effects of biogeochemical (carbon sequestration) and biogeophysical (energy balance) effects. These effects are known to vary at the continental scale (e.g., from boreal to tropical regions). Here, we show based on a four-year study that the biogeochemical vs. biogeophysical balance in paired forested and non-forested ecosystems across short distances and steep aridity gradient (∼200Km, aridity index 0.64 to 0.18) can change dramatically. The required time for the forestation cooling effects via carbon sequestration, to surpass its warming effects, associated with the forests reduced albedo and suppressed longwave radiation, decreased from >200 years in the driest sites to ∼70 years in the intermediate and ∼40 years in the wettest sites. Climate-related benefits of forestation, previously considered at large-spatial scales, should be considered at high-spatial resolutions in climate-change mitigation programs aimed at taking advantage of the vast non-forested dry regions.

**Teaser:** Climate-related effects of afforestation can vary between cooling and warming effects across 200 km.

## Introduction

Forests have complex interactions with the climate system through biogeochemical and biogeophysical processes, with implications for the Earth’s radiation balance ^1–5^. The growth of forests can have contrasting effects on climate: accumulation of large amounts of carbon mitigates the current enhancement of the greenhouse effect ^6^. In contrast, dark forest canopies are often characterized by low albedo, thereby increasing radiation absorption by adding heat to the surface ^3,7^. Under humid conditions, the albedo effects can be compensated for by the latent heat flux (LE), but in water-limited regions, most of the forest radiation load is dissipated as large sensible-heat flux (H). Variations in these contrasting effects of the carbon storage and the enhanced radiation absorption under dry conditions have not been characterized well at high spatial resolution.

Previous studies have indicated the warming effect of forestation activities in the boreal area due to snow-albedo feedbacks, where the decrease in albedo is significant during the long snow-cover periods ^8,9^. Recent studies have indicated similar but relatively weaker trends in temperate regions ^10^. Zhang *et al*. ^11^ showed a local cooling effect in temperate eastern USA, with a more significant cooling effect of reforestation in warmer sites. Forestation in the tropics is thought to be beneficial to the global climate ^12^, mainly because of the high productivity under high water availability conditions ^6^. Across regions and biomes, from tropical to temperate and from tropical to boreal, a large decrease in the benefits from forest carbon sequestration potential (the net ecosystem productivity; NEP) of 30% and 70%, respectively, is generally expected ^7^.

Most previous studies relying on simulations have pointed out changes in radiative forcing associated with forestation actions across biomes, with limited reference to small-scale variation ^1,13–15^. Such studies indicated that forestation actions, in which croplands or grasslands were converted to mature forests, reduced the carbon sequestration benefits due to changes in the surface radiation balance (e.g., reduced albedo). This reduced cooling effect was largest in the boreal region and decreased through the temperate zone to the tropical biomes, but the magnitude and direction of such effects can also change when additional factors are considered. For example, simulated afforestation in the tropics showed amplification of the biogeochemical cooling effects when variations in sensible and latent heat fluxes were considered ^15^. However, essentially all of these and similar studies rely on remote-sensing data and the sparsely distributed flux tower in the global network.

Drylands are defined as areas where the aridity index (AI, the ratio of annual precipitation to potential evapotranspiration) is below 0.65. These regions are commonly further divided into four aridity index categories ^16^: hyper-arid (AI≤0.05), arid (0.05<AI≤0.2), semi-arid (0.2<AI≤0.5), and dry-subhumid (0.5<AI≤0.65). Combined, drylands cover approximately 41% of the earth’s surface land area and 17 % of the global drylands are covered by forests^17–19^. Therefore, investigating processes that can help assess and evaluate the impact of changes in dryland forest characteristics are clearly important..

Hot drylands are subject to high solar radiation loads and low water availability. These distinctive conditions, in turn, may be reflected in the distinct climatic impacts of forestation in these regions. Dryland afforestation was found to have a strong warming effect due to the shortwave forcing associated with a decrease in albedo ^20^. Such effects can be further enhanced due to efficient canopy cooling through sensible heat (“convector effect” ^20,21^), and the resulting suppression of the thermal radiation flux ^20^. These effects ultimately result in an increased radiation load. In contrast, using coupled land-atmosphere models, Syktus & McAlpine ^22^ and Yosef *et al*. ^23^ suggested that large-scale restoration of savannas woodland ecosystems and forest ecosystems in Australia and the Sahel regions could lead to a local cooling effect through a chain of biogeophysical processes.

Despite water limitations, semi-arid ecosystems are known to play an important role in the global carbon cycle and climate, particularly in the interannual variability of the terrestrial carbon sinks ^24,25^. Lal ^26^ suggested that there is a significant carbon sequestration potential in soil organics of dryland ecosystems (up to 20 Pg C). In a recent study at a semiarid timberline, Qubaja *et al*. ^27^ suggested a large potential carbon sink due to afforestation in the semiarid soils, which was also associated with the extended soil carbon turnover time (almost 60 years in the top 1 m). The importance of drylands in assessing climate mitigation potential is further enhanced by the large proportion of the global land surface they occupy ^17^ and its restoration potential ^28^ (see also www.greatgreenwall.org).

This study was motivated by the potential for carbon sequestration in drylands, combined with the need to account for contrasting biogeophysical effects to obtain a more realistic assessment of the climatic benefits from forestation actions. We hypothesized that across the steep climatic gradient, variations in factors such as aridity, soil, and plant productivity, typical of drylands, could result in considerable changes in the climate change mitigation potential of forestation at fine scales. This could have important implications for forestation strategies in these areas.

## Results

We tested our hypothesis using a mobile lab to obtain observationally-based information on the radiative (long and shortwave radiation) and non-radiative (sensible heat, latent heat, and carbon) fluxes in paired Aleppo pine forest and non-forest sites across the steep climatic gradient in Israel (∼200km, from arid through semi-arid and dry-subhumid; Fig. 1, Table S1). Correlations between short-term campaigned based radiative and non-radiative fluxes and continuous meteorological data from nearby meteorological stations were combined to estimate the long-term radiative and non-radiative fluxes (using methodology previously developed; Rohatyn *et al*. ^29^ and see Methods for more information). The resulted fluxes were then averaged and summed over the period of 10-15 years and described here.

**Fig. 1.**
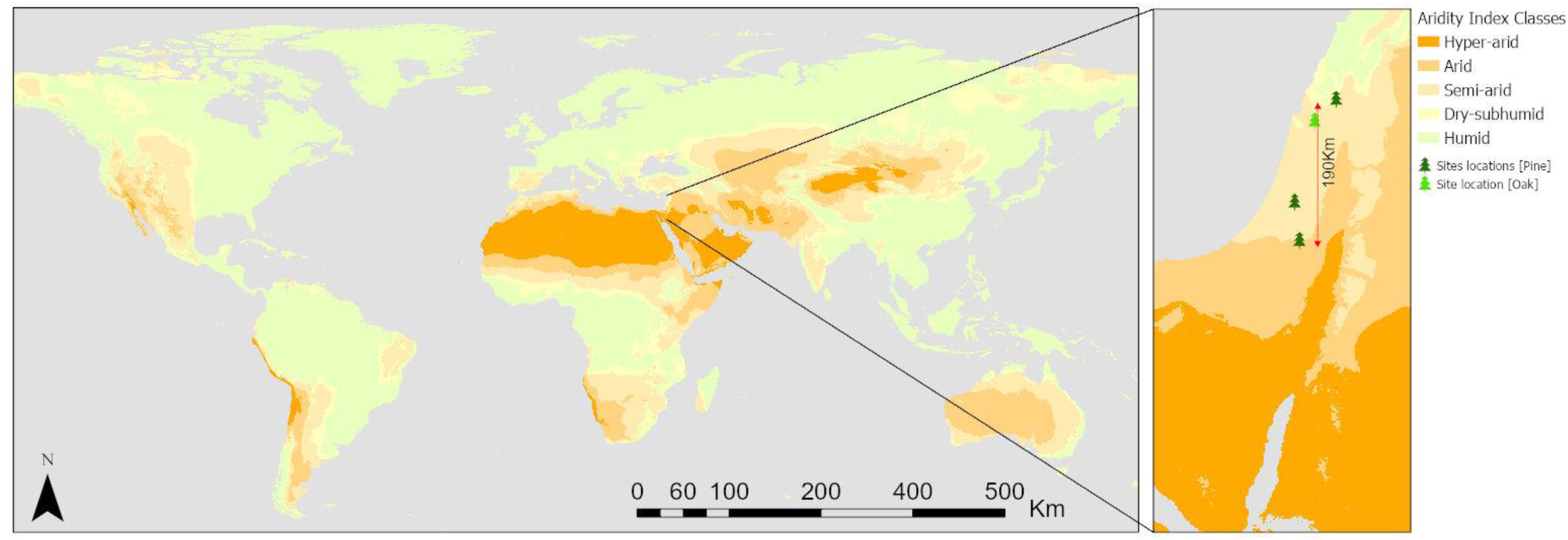
Global aridity index map and location of study sites across the climatic gradient in Israel. The Aridity Index was calculated as the ratio between Mean Annual Precipitation and Mean Annual Potential Evapotranspiration, according to Trabucco & Zomer, (2019; Global AI v2). AI was then used to classify hyper-arid (AI<0.05), arid (0.05<AI<0.2), semi-arid (0.2<AI<0.5), dry-subhumid (0.5<AI<0.65), and humid (AI>0.65) areas. The enlarged inset map shows the climatic gradient in Israel, indicating the paired sites locations of forested (pine or oak) and adjacent non-forested ecosystems. AI classes are presented with the color palette from orange (hyper-arid) to green (humid).

Afforestation (indicated here by the difference between forest and nearby non-forest sites) had significant effects on all measured components of surface-atmosphere exchange, as reflected in the observed, annual scale, changes in albedo, sensible heat, latent heat, and net radiation and carbon fluxes. At all forested sites, the albedo was significantly (P_value < 0.005) lower than that of the non-forested sites, which was essentially independent of the site (∼0.13 range across forested sites, which is ∼0.1 lower than the adjacent non-forested sites). At the non-forested sites, albedo increased across the climatic gradient (from 0.2 to 0.27; Fig. 2A), possibly associated with changes in soil properties (see Methods section and Table S1). Changes in radiation fluxes were also observed in the longwave components, with less negative values (i.e., reduced emission) of net longwave radiation (LWnet) in forested ecosystems, with the greatest effect in the arid sites (∼20 Wm^-2^; Fig. 2B). The combined effect of the reduced albedo and reduced longwave emissions contributed to the significant increase in net radiation in the forested sites (up to ∼50 Wm^-2^; Fig. 2D; P_value < 0.005).

**Fig. 2.**
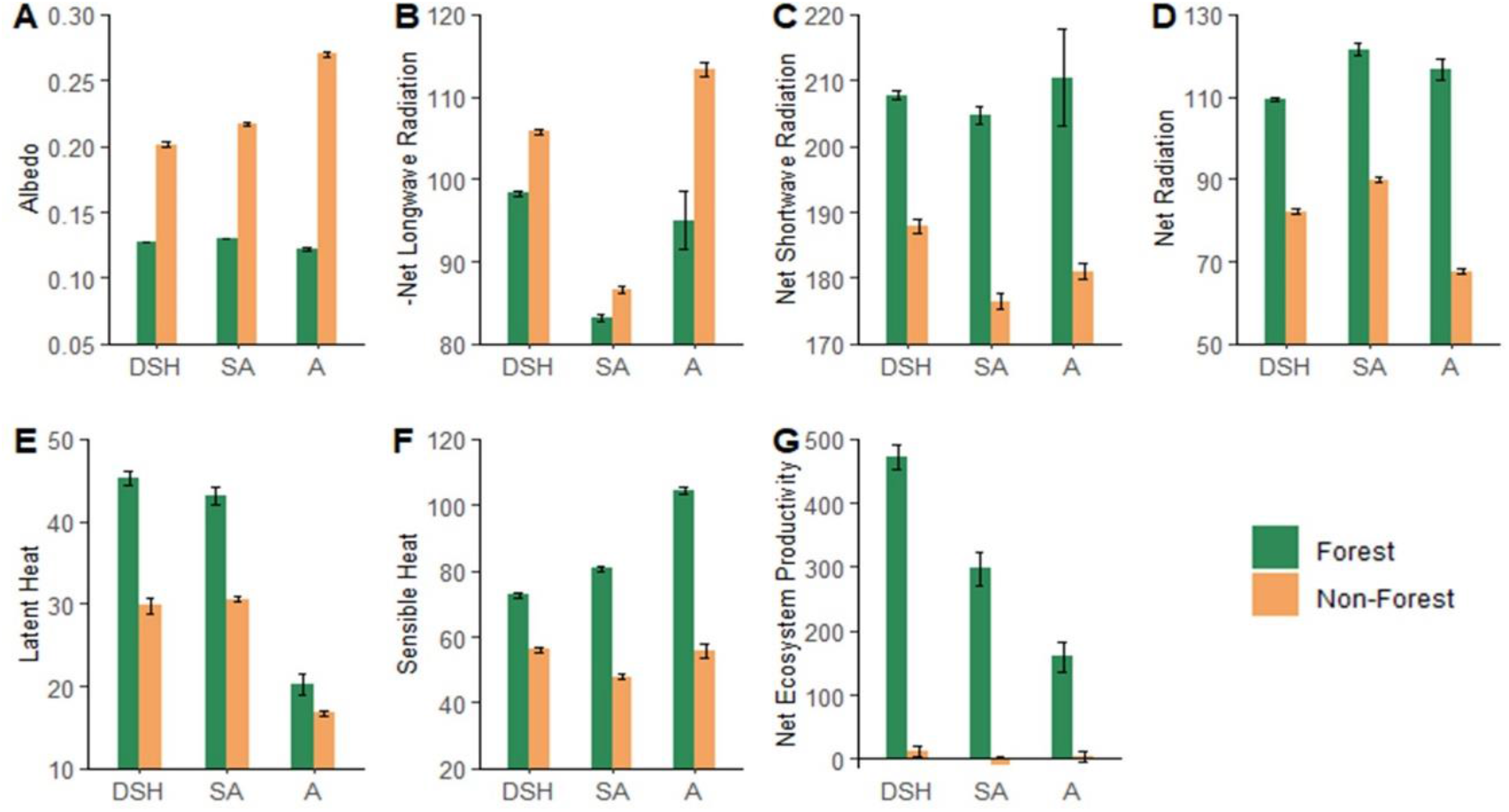
Forestation effects on the annual mean (2000–2015) values of the radiative budget components and net ecosystem production. Annual mean of Albedo (**A**), Net Shortwave Radiation (SWnet in W m^-2^; **B**), −Net Longwave Radiation (−LWnet in W m^−2^; **C**), Net Radiation (Rn in W m^−2^; **D**), Latent heat flux (LE in W m^−2^; **E**), Sensible heat flux (H in W m^−2^; **F**), and an annual sum of Net Ecosystem Productivity (NEP in gC m^−2^ yr^−1^; **G**). Values are from the forest (green bars) and non-forest (orange bars) ecosystems, across a climatic gradient, with dry-subhumid (DSH) to semi-arid (SA) and arid (A) climatic conditions. Note, the scaling differences among the sub-figures. The paired t-test was used to compare multi-annual averages of all variables between forested and adjacent non-forested sites and between sites across the climatic gradient. The error bars are for ± standard errors pf the means.

The higher net absorbed radiation in the forested sites was compensated for by higher non-radiative fluxes. As expected, the latent heat flux was always higher in the forested sites (50% and 40% higher in the more humid sites; P_value < 0.005), but the effect was minimal (only 20% higher; P_value = 0.007) in the arid site where essentially all annual precipitation is evaporated, independent of land cover (Fig. 2E). Sensible heat flux was always higher in the forested sites, but particularly in arid sites where it was the major energy outlet, which almost entirely balanced net radiation (>90%; Fig. 2D, F).

All non-forested sites had nearly zero net ecosystem productivity (NEP), which reflected the large contribution of annual vegetation with large seasonal fluxes of photosynthetic uptake and respiration, but negligible long-term carbon sequestration. In contrast, all forested sites had a significant NEP, consistent with the values of forest ages of approximately 40–50 years. NEP increased along the climatic gradient, with the dry-subhumid forested site showing almost three times higher NEP compared to that of the arid site (Fig. 2G).

To expand our study beyond the pine forest plantation we added measurements with the mobile lab in a native Oak-forest ecosystem. The comparison of the Oak-forest site to the Pine-forested and non-forested sites under the same climate (Fig. 3) indicated a lower albedo than the non-forested site (50% lower; P_value < 0.005), but only slightly higher value (10% higher; P_value < 0.005) than that of the Pine-forest ecosystems (Fig. 3A). Longwave radiation emission was reduced in the Oak-forest compared with that in the non-forested sites and to a greater extent than that in the Pine-forested site (Fig. 3B). Consistent with the albedo and LW radiation effects, the net radiation was markedly higher in the Oak-forest than in the non-forested sites, similar to the effect of the Pine forestation (Fig. 3D; P_value < 0.005). As in the pine forestation, latent heat fluxes increased to a similar extent in the Oak-forest site, compared with the non-forested site. However, the sensible heat flux was enhanced to a lesser extent in the Oak-forest than in the Pine-forest (Fig. 3E, F). Finally, while a significant NEP was observed in the Oak-forest, compared with the non-forested sites, it was nearly a third of the NEP in the Pine-forested sites (Fig. 3G; P_value < 0.005).

**Fig. 3.**
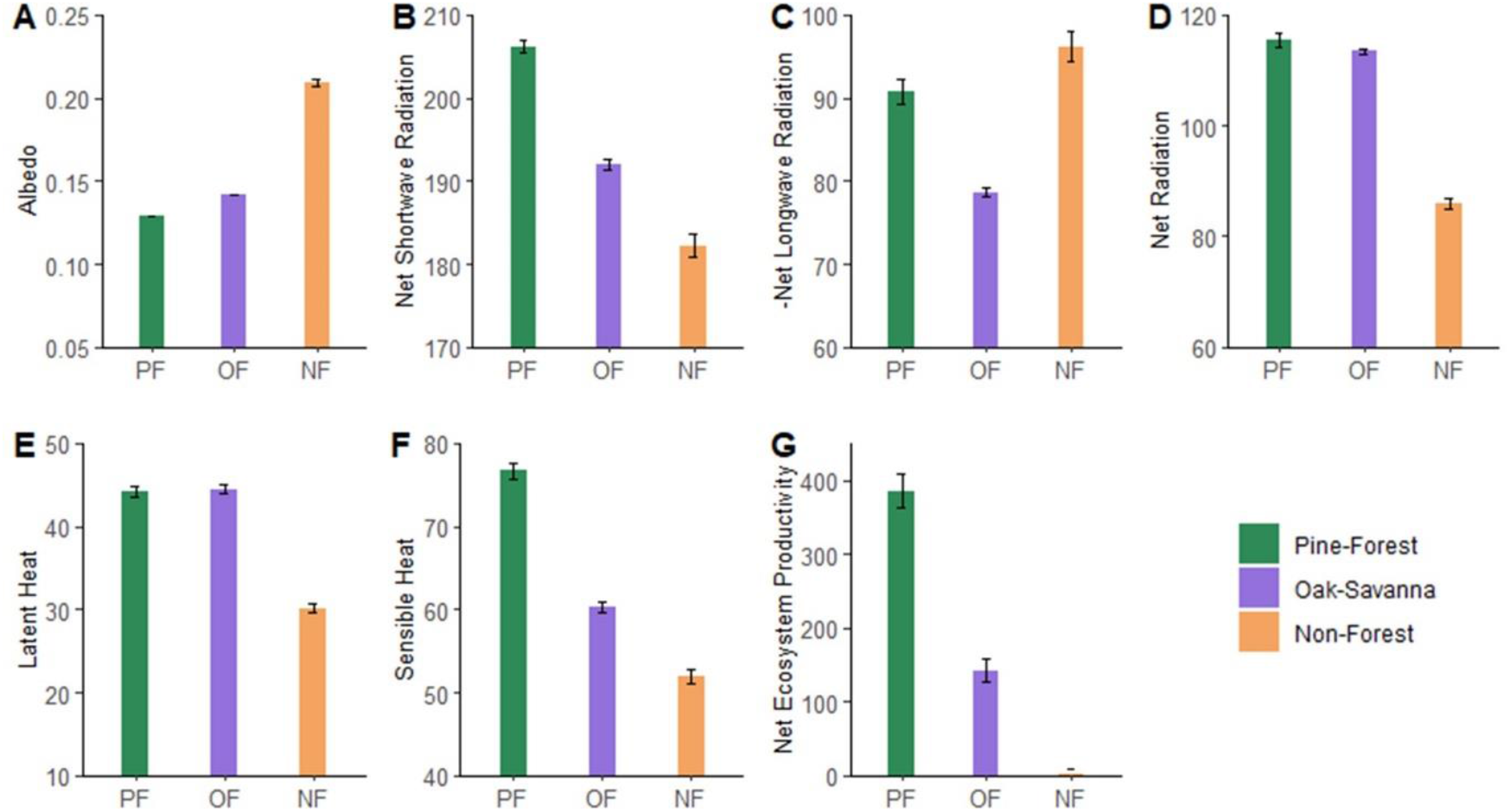
Land cover type effects on the annual means (2000–2015) of radiative and non-radiative fluxes. Annual mean of Albedo (**A**), Net Shortwave Radiation (SWnet in W m^-2^; **B**), −Net Longwave Radiation (−LWnet in W m^−2^; **C**), Net Radiation (Rn in W m^−2^; **D**), Latent heat flux (LE in W m^−2^; **E**), Sensible heat flux (H in W m^−2^; **F**), and an annual sum of Net Ecosystem Productivity (NEP in gC m^−2^ yr^−1^; **G**). Oak-forest (OF, purple bars) values compared with Aleppo Pine-forest (PF, green bars) and non-forest (NF, orange bars) averaged from multi-year annual means of the semi-arid and dry-subhumid paired sites. The paired t-test was used to compare multi-annual averages of all variables between forested and adjacent non-forested sites and between sites across the climatic gradient. The error bars are for ± standard errors pf the means.

The differences between forest and non-forest sites (Δ) are summarized in Table 1. Radiative changes (both changes in net shortwave, ΔSWnet, and net longwave, ΔLWnet, radiation) were compared with changes in carbon stocks (ΔNEP). As can be expected due to their close proximity, the incoming shortwave and longwave radiation for the paired sites were not significantly different (data not shown), and ΔSWnet and ΔLWnet reflect differences in the outgoing radiation. In all paired sites of Pine-forest vs. non-forest ecosystems, the impact of forestation on shortwave radiation fluxes (increase input) was larger than the effects on longwave radiation (suppressed output). These differences between forested and non-forested areas, in both increased input (ΔSWnet) and suppressed output (ΔLWnet), generally increased with drying across the climatic gradient (Table 1). However, in the case of the Oak-forest vs its paired non-forested sites, the differences in shortwave radiation were smaller than the differences in longwave radiation (4% and 8% of global radiation, Rg, respectively). Overall, ΔSWnet and ΔLWnet increased by 50% and 150%, respectively across the climatic gradient. Conversely, the differences in the annual net change in carbon stocks between forest and non-forest (ΔNEP; Table 1) decreased across the climatic gradient by 70%, from dry-subhumid to arid areas.

**Table 1.** Differences between forested (F) and adjacent non-forested (NF) for the three pine sites and one oak site in radiation budget and carbon stock. aridity index (AI), annual sums of precipitation (P, mm yr^−1^), annual means of global radiation (Rg in W m^−2^), annual minimum and annual maximum air temperature (Ta min-max in °C), annual means of changes (Δ=F−NF) in shortwave and longwave net radiation (ΔSWnet and ΔLWnet in W m^−2^), and changes in average annual sum (15 y) of net ecosystem productivity (NEP in gC m^−2^ yr^−1^). The last three columns present the years needed to balance between the warming effect due to SW and LW forcing by the cooling effect of carbon stock change (‘Break even time’ in years) according to Eq. 7.

| | AI<br>(#) | P<br>(mm<br>yr<sup>-1</sup>) | Rg<br>(W m<sup>-2</sup>) | Ta<br>min-max<br>(°C) | ΔSWnet<br>(W m<sup>-2</sup>) | ΔLWnet<br>(W m<sup>-2</sup>) | ΔNEP<br>(gC m<sup>-2</sup><br>yr<sup>-1</sup>) | ‘Break even time’ | | |
| --- | --- | --- | --- | --- | --- | --- | --- | --- | --- | --- |
|  |  |  |  |  |  |  |  | SW<br>forcing<br>(years) | LW<br>forcing<br>(years) | Total<br>(years) |
| **Dry-Subhumid** | 0.64 | 770 | 235 | 5–27 | 19.8 | 7.5 | 460 | 31 | 12 | 43 |
| **Semi-arid** | 0.37 | 480 | 228 | 11–30 | 28.2 | 3.4 | 310 | 65 | 8 | 73 |
| **Arid** | 0.18 | 280 | 242 | 6–28 | 29.4 | 18.3 | 160 | 132 | 81 | 213 |
| **Oak-forest** | 0.4 | 580 | 223 | 9–29 | 9.8 | 17.6 | 142 | 42 | 89 | 131 |

We then used a common method to compare radiation and carbon fluxes by converting the net change in carbon flux due to forestation action (forest minus non-forest) to the resulted change in radiative forcing based on the CO_2_ radiative efficiency (Myhre *et al*. ^30^). Combined, the radiation and carbon components indicated a sharp increase across the climatic gradient in the ‘Break-even time’, the number of years of carbon accumulation that produce the radiative forcing required to balance that of the change in albedo and LW radiation suppression (Table 1). In the dry-subhumid sites, the ‘Break-even time’ was 43 years (31 years based on the shortwave and 12 years based on the longwave components). In the arid sites, the time to balance was 213 years (132 years for shortwave and 81 for longwave; note that this >200 years computation result has no special significance beyond indicating that it is well beyond the relevant time horizon for carbon accumulation in dry forests). For the Oak-forest, the ‘Break-even time’ was 42 years based on the shortwave part, which was intermediate between the wetter pine sites. But it was 89 years based on the longwave component, which was longer than in all pine sites, resulting in a total ‘Break-even time’ of up to 131 years at this site.

Finally, we extrapolated our results from the different forest study sites to a single 80 years forest age basis and used the emission equivalent method of converting radiation fluxes into carbon equivalent units (see Methods eq. 8+9), to allow comparison of our small spatial scale (∼200 km) climatic gradient study with the results of four large-scale studies found to be sufficiently suitable for such comparison (Table 2). The results in three of the four studies indicated a larger net sequestration potential (ΔSP) in the temperate and tropical biomes than in the boreal region (approximately 50%–150% increase). Arora & Montenegro (2011) reported minimal differences in ΔSP between boreal and temperate biomes, but with a much higher ΔSP in the tropical biome (∼60% higher than temperate and boreal). Our ΔSP measurements, associated with the conversion of the non-forested sites to Pine-forests, indicated a similar magnitude of change, but across a much smaller spatial scale. Compared with the arid site, ΔSP increased by 85% in the semi-arid site and by almost 200% in the dry-subhumid site. Note that the non-forested SP for the current study is near zero at the annual scale (Fig. 2G) and therefore ΔSP in such cases provide an approximation for SP itself in the different sites, indicating a range between 120 and 220 tC ha^-1^ for 80 years of the forest growth (Table 2). This translates to 1.5 to 2.8 tC ha^-1^ yr^-1^, which is consistent within the range obtained by the global FLUXNET network^31^.

**Table 2.**
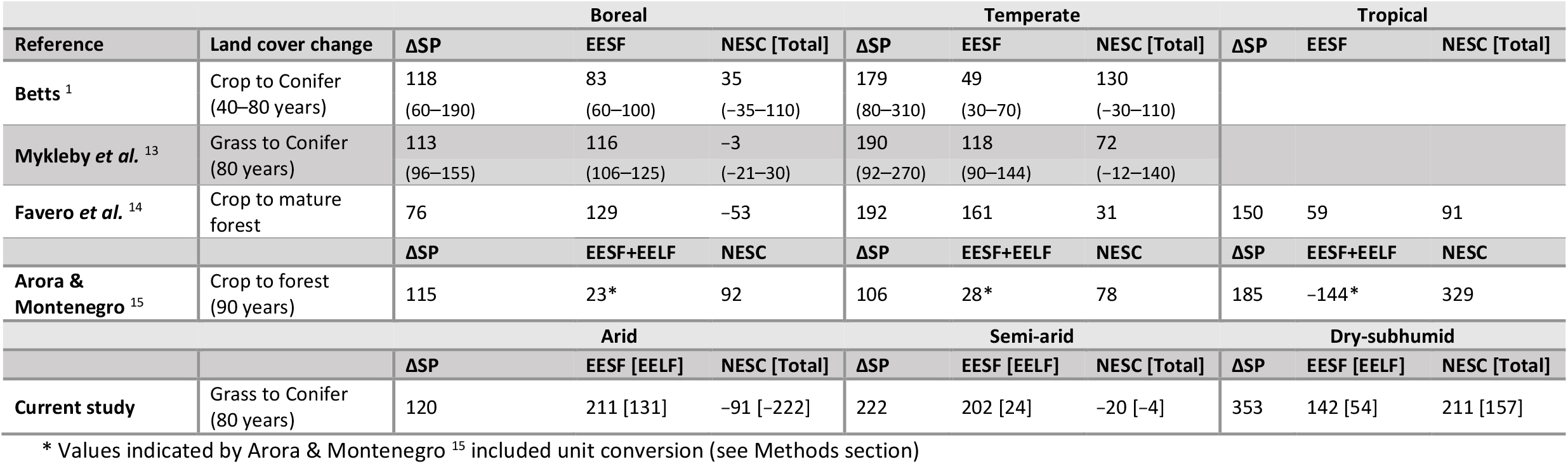
Comparison of global and large-scale regional studies with the current local study for the land cover changes effects on the surface radiative forcing. net sequestration potential (ΔSP, Eq. 9), emission equivalent of shortwave forcing (EESF, Eq. 10), and net equivalent stock change (NESC= SP−EESF, Eq. 11) are summarized based on published values from three large scale regional studies across three biomes (Boreal, Temperate, and Tropical). All values are in tC ha^−1^ accumulated over forest age or rotation period as specified in the second column brackets. The last two rows represent the current study extrapolated over 80 years since forest establishment for arid and semi-arid dry-subhumid forests. When the longwave radiation forcing was included in the analysis, +EELF is indicated for the combined effect of shortwave and longwave radiation forcing or [EELF] if the longwave is calculated separately.

The results for the estimations of the emission equivalent of shortwave forcing (EESF) differed in sign and magnitude between the different studies (Table 2). The study by Betts ^1^ was the only one to report lower EESF in the temperate compared with the boreal biomes (40% decrease). The study by Mykleby *et al*. ^13^ found almost no difference between the two regions, but with 2-3 times higher EESF compared to Betts ^1^. The studies of Favero *et al*. ^14^ and Arora & Montenegro ^15^ showed similar results for the EESF estimated values, with approximately 20–25% higher EESF in temperate biomes than in boreal biomes. In the last two studies, the EESF estimates were different in sign and magnitude, with positive but rather small EESF in the Favero *et al*. ^14^ study, and negative EESF indicated by Arora & Montenegro (2011). In comparison, in the present study, the EESF estimates were much higher in the arid and semi-arid sites than in the dry-subhumid climatic zones but, notably, with variations of the same order of magnitude as in the large-scale studies (Table 2). In our study, we also included the analysis of longwave forcing effects (EELF), which can be significant (e.g., Rotenberg & Yakir ^20^), and indicated a similar magnitude of spatial variability (Table 2).

## Discussion

### Forestation effect on the partitioning of radiative and non-radiative fluxes

The paired sites approach used here to compare forested vs. adjacent non-forest sites (i.e., sites under similar conditions) is useful for evaluating the projected outcomes of forestation and land-use changes in general ^32^. In this study, we also considered the effect of variations in climate on such projections by studying several paired sites across a sharp climatic gradient (in precipitation and, hence, in AI), and we focused on the dry Mediterranean region, which has significant potential for afforestation ^28^. Based on this approach we could obtain quantitative information that is critical in developing forestation and afforestation strategies for climate change mitigation that are often proposed for the vast dry-land regions ^33–35^.

As expected, the results demonstrated how forestation actions consistently decreased surface albedo, thus contributing to an increase in the net shortwave radiation at the surface. However, albedo change increased with increasing aridity. These changes across the climatic gradient resulted mainly from the characteristics of the non-forested sites, particularly vegetation cover and soil types. Vegetation cover is determined by climate, and the vegetation cover is of lower stature in the arid non-forested site than in the other sites. However, it is also greatly affected by grazing, which was, to our knowledge, more intense in the arid and dry-subhumid sites. The increase in albedo with increasing aridity in the non-forested sites could therefore be influenced by changes in the proportion of vegetation cover, which in fact increased the effects of changes in the soil type along the gradient. Soil type, in turn, is related to geographical distribution across the region. The soil type in the arid site (Rendzic Leptosol, or Haploxeroll) is brighter than the increasing contribution of the Terra Rossa soil type in the wetter sites ^36^. Notably, our result for the annual mean non-forest albedo in the arid site was 0.27, which is lower than the 0.3–0.35 range measured in the region only in summer by Ben-Gai *et al*. ^37^, but consistent with our summer value of 0.33. This demonstrates the importance of accounting for seasonal changes and may reflect variations in the combined effects of vegetation and soil. While the change in albedo varied along the aridity gradient, the albedo for the forested sites was essentially similar (Fig. 2). This is indicated by the minor changes in leaf properties in the Aleppo pine trees and little sensitivity to variations in tree density (see Methods section), partly because tree cover tends to be near-constant, compensating for stand density, and both measurements and sun angles are mostly off the nadir. It is likely that in contrast to the non-forested sites, the forest albedo in evergreen forests (but possibly not in deciduous forests) is essentially decoupled from changes in soil type.

The observed increase in net radiation along the gradient values in the forested sites (composed of the combined albedo and longwave radiation suppression effects) was compensated for by the increase in the non-radiative fluxes of sensible and latent heat. This is a natural mechanism for ecosystem energy balance management ^32^. However, the partitioning between the two fluxes differed along the aridity gradient. As expected, latent heat fluxes were high in the more humid sites, and sensible heat flux was high in the arid site. These differences are due to differences in available water for the evapotranspiration flux. However, the large sensible heat fluxes in the dry site also require low aerodynamic resistance, which is associated with the roughness of the more sparse canopy structure under these conditions (the generation of the “canopy convector effect”; ^20,21^) Interestingly, we have recently shown that convector-type non-evaporative canopy cooling may originate at the leaf scale ^38^. Irrespective of the mechanism, the more efficient canopy cooling, and the lower forest canopy (skin) temperature, in comparison with the non-forested land surface, feedback on their radiative fluxes by reducing the outgoing longwave radiation, compared with the non-forested area. The resulting increase in the net longwave radiation significantly amplifies the forest surface warming of the albedo effects. The processes discussed above are driven by moisture limitation and are therefore particularly pronounced in dry regions, and as shown here, considerably increase along the aridity gradient (Fig. 2). The low net longwave radiation in both semi-arid sites (Fig. 2B) and, in turn, the small changes in the longwave component (Table 1), may indicate some local warming effects, which have not been sufficiently identified at present.

### Forest species composition

While this study focused on comparing paired sites of maximum similarity and therefore used mainly Aleppo pine sites, that were common in the study area, an Oak-forest site nearby offered the opportunity to examine the effects of the dominant tree species. The nearby Oak-forest and Pine-forest sites had similar aridity but differed largely in vegetation characteristics and canopy structure. Nevertheless, the two forest types revealed similar changes in albedo relative to the non-forested sites. However, they markedly differed in the change in the net longwave radiation between the forest and non-forest sites (Table 1). Furthermore, while this ecosystem had larger sensible heat flux than the non-forested site, it was much smaller than in the pines, indicating a more limited ‘convector effect’ of this canopy. The suppressed LW radiation (cooler surface) and limited H, could be balanced by the relatively large latent heat flux associated with the more extensive inter-canopy grasses and annuals at this site. This difference can be due to the greater exposure of the soil surface at this site (smaller canopies). Note that the net ecosystem productivity (NEP) in the Oak-forest ecosystem was approximately one-third of that in the Pine-forest ecosystem. The low NEP can be associated with the smaller trees and tree canopies, which consists mostly of deciduous trees, exposing the grasses, and leading to high autumn-respiration rates, as was seen at the seasonal scale measurements (data not shown). Note that in this ecosystem, the balance of the net radiation by the LE + H fluxes seems to be lower than obtain for the pines (Fig. 3 D-F), but within the acceptable range (energy closure of approximately 0.9).

### ‘Break-even time’

The differences in radiative and non-radiative fluxes described above (Table 2) determine the biogeophysical effect of forestation action on radiative forcing. Comparing forest and non-forest paired sites showed in all cases that the forests’ biogeophysical effects produce “warming” effects, due to both shortwave and longwave forcing. This is generally a one-time change that can be expected to occur in the early stages of forestation ^20,39^. This warming effect could then be compared with the cumulative biogeochemical “cooling” effect due to forest carbon sequestration, which reduces the radiative forcing by reducing atmospheric CO_2_ concentrations. NEP increased significantly with decreasing aridity, except for the Oak-forest ecosystem, which had a much lower NEP, compared with all pine ecosystems. The high NEP at the dry-subhumid site (460 gC m^-2^ yr^-1^) is in agreement with unpublished data for a Greek flux site at Sani, with similar climate conditions, and within the range reported for Mediterranean warm evergreen forests (380 ± 73 gC m^-2^ yr^-1^)^31^. The combination of high NEP and relatively low shortwave and longwave forcing in the dry-subhumid site resulted in the lowest estimation of the ‘Break-even time’, of 43 years (Table 1), which is already below the current age of this still highly productive forest.

In contrast to the dry-subhumid site, the low NEP and high shortwave and longwave forcing at the arid site resulted in a very high estimated ‘break-even time’ of 213 years, which is possibly longer than the forest life cycle. This long ‘break-even time’ is at least partially due to the low vegetation cover, based on an annual average, in the paired non-forested site. This, in turn, is the combined result of the dry conditions and intensive grazing and could enhance desertification processes, as shown for semi-arid lands in Inner Mongolia ^40^. In contrast, some ameliorating effects, such as taking advantage of the topography and limiting forestation to northern slopes, which could reduce the albedo effects and enhance productivity ^41,42^, also deserve further research. Accordingly, we suggest that in considering afforestation actions in drylands as a climate mitigation tool, their actual climatic benefits should be first examined at the local scale. This is the case in particular, in areas with characteristics (vegetation cover and soil properties), such as in the current arid study site, where afforestation could result in climatic effects, which are in contrast to initial expectations.

The importance of the dominant tree species in the forestation system was apparent in the comparison of the Oak-forest and Pine-forest sites to the respective non-forest sites. While the oaks and pines had relatively similar effects on net radiation, the low NEP in the Oak-forest resulted in a much longer ‘break-even time’ (Fig. 3, Table 1). The Oak-forest ecosystem is a typical and relatively common ecosystem for the study area ^43^, and the ‘break-even time’ analysis (large biogeophysical and low biogeochemical effects) makes the climatic benefits of this ecosystem type questionable as the leading motivation for its expansion. However, the planted Pine-forests (using Aleppo pines, which also have native history; ^44^) showed much higher NEP, and shorter ‘break-even time’. Such more favorable balance can also be expected for the similarly common but more dense Mediterranean woodlands, which include oaks as one component ^45^. Furthermore, these ecosystems are important for preserving biodiversity and a range of other ecological services in this region (e.g., wood production, erosion protection, and leisure activities, among other services; ^46^), irrespective of their contribution to the global climate. Note, however, that the results from the Oak-forest are based on one site, and future expansion of studies on the climatic benefits of this locally important ecosystem is strongly encouraged.

Our results indicate that differences in conditions (aridity and soil) in regions exposed to similarly high solar radiation (annual mean of 220–240 W m^−2^), and even within small geographic distances, may amplify the biogeophysical effects of forestation that have a warming effect ^47^. The results demonstrate that the forest effect on both shortwave and longwave forcing can be much larger in semi-arid and arid regions than in more humid regions. It also demonstrates the sensitivity of the climatic effects of forestation actions to changes over a small spatial scale in the dry-subhumid to arid transition zone. Further, they indicate that the benefits of afforestation for climate change mitigation diminish with increasing aridity.

It has recently been indicated ^19^ that under the large-scale drying process expected under climatic changes in a business-as-usual scenario, 11.2% of the land area will shift to a drier class of aridity by the end of the century. A cross-analysis of Koutroulis ^19^ results and the map of potential restoration opportunities of the Global Restoration Initiative (GRI;^48^), indicated that ∼450 Mha of land with a potential of restoration will shift to a drier aridity class by 2100. Since the ‘break-even time’ increases with aridity, and may exceed the expected forests life cycles, our results indicate that shifts such as predicted by Koutroulis ^19^ may result in corresponding shifts in the potential climate change mitigation benefits of forestation actions in drylands. This is also applicable to various proposed natural-based solution activities in the vast dryland regions (∼ 40% of the Earth’s land surface area; ^49^), especially those under high radiation load. Importantly, however, forests support other ecosystem services, apart from the direct mitigation of global warming. Large-scale dryland afforestation can also modify local and regional precipitation and surface temperature, such as recently demonstrated for Sahel and Australia ^23^. Similarly, in a regional study in Australia, savanna restoration increased local biophysical cooling ^22^. Finally, it is important to note that focusing on the ‘break-even time’ metrics is not an alternative to considering the potential for carbon sequestration by forests. Carbon sequestration is assessed by a variety of metrics, such as stem carbon (e.g. Braakekke *et al*. ^50^), timber and biomass, and wood production (e.g. Anderson *et al*. ^51^; Favero *et al*. ^14^), or focus on the rate of accumulation using growth curves (e.g. Nilson and Peterson ^52^; Braakekke *et al*. ^50^). Here we use SP in tC ha^-1^, which can be interconverted with these metrics. However, focusing on the ‘break-even time’ analysis as done here, is critical even before providing an assessment of the penalty associated with factors such as albedo change, which we show can greatly increase in semi-arid conditions annulling the sequestration issue altogether.

### Forest climatic benefit - from continental scale to short distances

Our results (Table 2) reveal how climatic benefits from forestation actions can vary significantly across short distances along a climatic gradient (<200 km). Such variations, and in the same order of magnitude, have been previously reported only across large distances, typically at continental scales. Furthermore, while biome-scale considerations are often made, such as in Griscom *et al*. ^12^, the potentially large variabilities at finer scales, such as the inclusion of the importance of species composition, soil and ecosystem types as shown here are still rarely considered. Therefore, our results demonstrate the importance of high-resolution studies, especially when considering climate change mitigation strategies that focus on taking advantage of the forestation potential of the vast dry land area. We suggest that afforestation of drylands ecosystems where long ‘break-even time’ is expected (sometimes well beyond the relevant time scale for forest carbon sequestration), should be avoided as means of climate change mitigation. Nevertheless, the afforestation of such lands could remain relevant due to other potential ecosystem services, such as combating desertification, prevention of soil erosion, local wood production, and social aspects like shading and recreation).

## Materials and Methods

### Study site description

The study was carried out at the edge of an arid region in mature plantations dominated by *P. halepensis*, of an age of 40-50 years (*Pinus halepensis*), and their adjacent non-forested ecosystems. The sites were distributed across a climatic gradient from arid and semi-arid to dry sub-humid (Figure 1, Table S1). Three selected paired sites included: (1) An arid site at Yatir forest (annual precipitation, P: 280 mm; aridity index, AI: 0.18; elevation: 650 m; light brown Rendzina soil, and forest density: 300 trees ha^−1^), where a permanent flux tower has been operating since the year 2000 (http://fluxnet.ornl.gov). Note that an AI of 0.2 marks the boundary between arid and semi-arid regions. Yatir, with AI=0.18, is formally within an arid zone, but on the edge of a semi-arid zone. (2) An intermediate semi-arid site in Eshtaol forest (P: 480 mm; AI: 0.37; elevation: 350 m; light brown Rendzina soil, and forest density: 450 trees ha^−1^). (3) A dry-subhumid site in northern Israel at the Birya forest (P: 770 mm; AI: 0.64; elevation: 755 m; dark brown Terra-Rossa and Rendzina soil, and forest density: 600 trees ha^−1^). Non-forest ecosystems were sparse dwarf shrublands, dominated by *Sarcopoterium spinosum* in a patchy distribution with a wide variety of herbaceous species, mostly annuals, that grew in between the shrubs during winter to early spring, and then dried out. All shrubland sites had been subjected to livestock grazing (exposing soils). Finally, an additional site that was characterized as Oak-forest vegetation was added. The site was dominated by two oak species, *Quercus calliprinos* and *Quercus ithaburensis* (P: 540 mm; AI: 0.4; for more details on the oak site, refer to Llusia *et al*. ^53^). All sites were under high solar radiation load, with an annual average of approximately 240 Wm^-2^ in the arid region and only 3% lower in the northern site in the dry-subhumid region (Table 1).

### Mobile laboratory

Measurements were conducted on a campaign basis using a mobile lab with a flux measurement system at all sites except the Yatir forest, where the permanent flux tower was used (http://fluxnet.ornl.gov; ^54^). Repeated campaigns of approximately two weeks at each site, along the seasonal cycle, were undertaken during 4 years of measurements, 2012–2015 (a total of 6-7 campaigns per site, evenly distributed between the seasons) the 4 years of measurements were found to be representative of previous 70 years of precipitation record (Figure S1). The mobile lab was housed on a 12-ton 4 × 4 truck with a pneumatic mast with an eddy-covariance system and provided the facility for any auxiliary and related measurements. Non-radiative flux measurements were undertaken using an eddy-covariance system to quantify CO_2_, and sensible heat (H), and latent heat (LE) fluxes using a 3D sonic anemometer (R3, Gill Instruments, Hampshire, UK) and an enclosed-path CO_2_/H_2_O infrared gas analyzer (IRGA; LI-7200, LI-COR). Non-radiative flux measurements were accompanied by meteorological sensors, including air temperature (Ta), relative humidity (RH), and pressure (Campbell Scientific Inc., Logan, UT, USA), radiation fluxes of net solar- and net long-wave radiation (SWnet and LWnet, respectively), and photosynthetic radiation sensors (Kipp & Zonen, Delft, Holland). Raw EC data and the data from the meteorological sensors were collected using a computer and a CR3000 logger (Campbell Sci., Logan, UT, USA), respectively. The EC system was positioned at the center of each field site with the location and height aimed at providing sufficient ‘fetch’ of relatively homogeneous terrains. For detailed information on the use of the mobile lab and the following data processing of short and long-term fluxes see Asaf *et al*. and Rohatyn *et al*. previous publications ^29,55,56^.

### Data processing

Mean 30-min fluxes (CO_2_, LE, and H) were computed using Eddy-pro 5.1.1 software (LiCor, Lincoln, Nebraska, USA). Quality control of the data included a spike removal procedure. A linear fit was used for filling short gaps (below three hours) of missing values due to technical failure. Information about background meteorological parameters, including P, Ta, RH, and global radiation (Rg), was collected from meteorological stations (standard met stations maintained by the Israel Meteorological Service, https://ims.data.gov.il/). The data were obtained at half-hourly time resolution and for a continuous period of 15 years since 2000.

### Estimating continuous fluxes using the flux meteorological algorithm

Estimation of the flux-based annual carbon and radiation budgets was undertaken using the short campaign measurements as a basis to produce a continuous, seasonal, annual, and inter-annual scale dataset of ecosystem fluxes (flux meteorological algorithm). The flux meteorological algorithm method was undertaken based on the relationships between measured fluxes (CO_2_, LE, H, SWnet, and LWnet) and meteorological parameters (Ta, RH, Rg, VPD, and transpiration deficit, a parameter that correlated well with soil moisture, see main text and supplementary material of Rohatyn *et al*. ^29^. A two-step multiple stepwise regression was established, first between the measured fluxes (H, LE, and the ecosystem net carbon exchange) and the meteorological parameters measured by the mobile lab devices, and then between the two meteorological datasets (i.e., the variables measured by the Israel meteorological stations) for the same measurement times. Annual fluxes were calculated for the combined dataset of all campaigns at each site using the following generic linear equation:

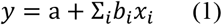

where, y is the ecosystem flux of interest, the daily average for radiative fluxes (LWnet and SWnet), non-radiative fluxes (H and LE), and daily sum for net ecosystem exchange (NEE), a and *b*_*i*_ are parameters, and *x*_*i*_ is Ta, RH, Rg, vapor pressure deficit, or transpiration deficit. The meteorological variables (*x*_*i*_) were selected by stepwise regression, with *b*_*i*_ = 0 when a specific *x*_*i*_ was excluded.

Based on this methodology, ecosystem flux data were extrapolated to the previous 7–15 years (since 2000 in the dry-subhumid and arid sites, since 2004 in the semi-arid sites, and since 2008 in the Oak-forest site) using all the available continuous meteorological parameters from the meteorological stations associated with our field sites. The long-term annual sums and means of extrapolated ecosystem fluxes were averaged for multi-year means of each site for the period of available extrapolated data. To test the extrapolation method, we conducted simulation experiments at the arid forest site, where continuous flux data from the 20 years old permanent flux tower were available. Five percent of the daily data were selected by bootstrap, a stepwise regression was performed for this sample, and then, the prediction of fluxes using the eq. 1 above was performed for the entire observation period (R^2^ of about 60% for the NEE flux; see Rohatyn *et al*. ^29^ for more details).

The aridity index of the Oak-forest was in between those of the semi-arid and dry-subhumid Pine-forests (0.4 compared to 0.37 and 0.67, respectively). Therefore, to compare the Oak-forest with Pine-forest and non-forest sites, the average results from the semi-arid and dry-subhumid paired sites were used.

### Radiative Forcing and Carbon Equivalence Equations

To compare the changes in the carbon and radiation budgets caused by forestation, we adopted the approach of Myhere *et al*. ^30^, and used Eq. 2:

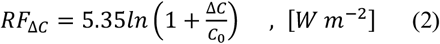

where land-use changes in radiative forcing (*RF*_Δ*C*_) are calculated based on the CO_2_ reference concentration, C_0_ (400 ppm for the measured period of study), and Δ*C*, which is the change in atmospheric CO_2_ in ppm, with a constant radiative efficiency (RE) value of 5.35. Here, Δ*C* is calculated based on the annual net ecosystem productivity (NEP; positive carbon gain by the forest, which is identical to net ecosystem exchange (NEE), the negative carbon removal from the atmosphere) as the difference between forested and non-forested ecosystems (ΔNEP) multiplied by a unit conversion constant:

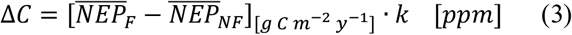

where, k is a unit conversion factor, from ppm to g C (k =2.13 × 10^9^), calculated as the ratio between the air molar mass (M_a_= 28.95; g mol^−1^), to carbon molar mass (M_c_= 12.0107; g mol^−1^), and total air mass (m_a_=5.15 × 10^9^; g).

Etminan *et al*. ^57^ introduced an updated approach to calculate the RE as a co-dependent of the change in CO_2_ concentration and atmospheric N_2_O:

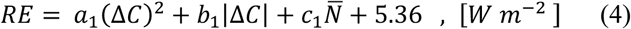

where, Δ*C* is the change in atmospheric CO_2_ in ppm resulting from the forestation, as calculated in Eq. 3, 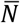 is the atmospheric N_2_O concentration in ppb (323), and the coefficients a_1_, b_1_, and c_1_ are −2.4 × 10^−7^ Wm^−2^ppm^−1^, 7.2 × 10^−4^ Wm^−2^ppm^−1^, and −2.1 × 10^−4^ Wm^−2^ppb^−1^, respectively.

Combining Eqs. 2 and 4 with an airborne fraction of *ζ* = 0.44 ^58^, we obtain Eq. 5:

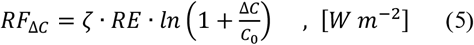

Next, the annual average radiative forcing due to differences in radiation flux was calculated as follows:

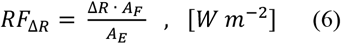

where, Δ*R* is the difference between forest and non-forest reflected short-wave or emitted long-wave radiation (Δ*SWnet* and Δ*LWnet*, respectively), assuming that the atmospheric incoming solar and thermal radiation fluxes are identical for the two, normalized by the ratio of the forest area (*A*_*F*_) to the Earth area (*A*_*E*_ = 5.1 × 10^14^ *m*^2^).

As forest conversion mostly has a lower albedo, the number of years needed to balance (‘Break even time’) the warming effect of changes in radiation budget by the cooling effect of carbon sequestration is calculated by combining Eqs. 5 and 6:

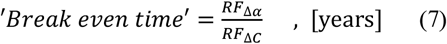

The multiyear averages of NEP for each of the three paired sites (forest and non-forest) were then modeled over a forest life span of 80 years. This was done based on a logarithmic model, modified for dryland, which takes as an input the long-term averages of NEP 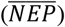:

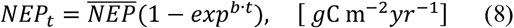

where annual carbon gain at time t (*NEP*_*t*_) is a function of the multiyear average carbon gain 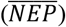, forest age (*t*), and growth rate (*b*). Parameter b is a constant (b = −0.17) and is calculated based on the global analysis of Besnard *et al*. ^59^, limiting the data to only dryland flux sites ^60^. Note that the analysis based on Besnard *et al*. ^59^ indicates NEP reaching a steady state and was used here to describe the initial forest growth phase, while growth analyses indicate that carbon sequestration peaks after about 80 years, followed by a steep decline ^50^. This is consistent with the time scale for forest carbon sequestration considered here.

The net sequestration potential (ΔSP) for each of the paired sites was calculated as the accumulated ecosystem Δ*NEP*_*t*_ along with forest age (Δ is the difference between forest and non-forest sites):

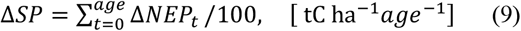

The ΔSP growth model was compared with previously published data of long-term carbon stock changes in arid forests (i.e., cumulative NEP over 50 years since forest establishment, t = 50), demonstrating agreement within ±10% ^27^.

For comparison with previous studies, the carbon emission equivalent of shortwave forcing (EESF) was calculated using an inverse version of Eqs. 5 and 6 based on Betts ^1^:

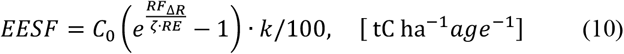

where, C_0_ is the reference atmospheric CO_2_ concentration (400 ppm, the average atmospheric concentration for the past decade), *RF*_Δ*R*_ is the multiyear average change in radiative forcing as a result of the change in surface albedo (Eq. 6 W m^−2^), *RE* is the radiative efficiency (Eq. 4, W m^−2^), ζ is the airborne fraction (0.44 as in Eq. 5), and k is a conversion factor, from ppm to g C (2.13 × 10^9^ as in Eq. 3). Eq. 10 was also used to calculate the emission equivalent of longwave forcing (EELF) with the *RF*_Δ*R*_ as the multiyear average change in radiative forcing as a result of the change in net long-wave radiation (Δ*LWnet*).

Finally, the net equivalent change in carbon stock due to both the cooling effect of carbon sequestration and the warming effect due to albedo change (net equivalent stock change; NESC), was calculated by a simple subtraction:

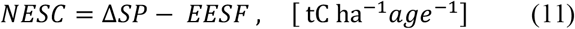

A comparison of the ΔSP (Eq. 9), the EESF (Eq. 10), and NESC (Eq. 11) with the same metrics as those used in other studies ^1,14,61^ was done when appropriate. An exception was made for Arora & Montenegro (2011), where only carbon stock changes (ΔSP) were available in carbon units, and biogeophysical (BGP) and biogeochemical (BGC) effects were expressed as temperature changes. To overcome this metric difference, we converted the biogeophysical to carbon equivalent units (EESF+EELF) by multiplying the carbon stock changes (ΔSP) by the ratio between the BGP and BGC effects on temperature (EESF+ EELF= ΔSP × BGP/BGC).

### Statistical and data analyses

The paired t-test was used to compare multi-annual averages of all variables between forested and adjacent non-forested sites and between sites across the climatic gradient. The variables of interest that were detected for their significant differences were albedo, net radiation and its longwave and shortwave components, latent heat fluxes, sensible heat fluxes, and net ecosystem productivity. All statistical and data analyses were performed using R 3.6.0 (R Core Team, 2020).

## Acknowledgments

S.R., E.R, F.T., Y.C., and D.Y. jointly planned and designed the methods and results of the research. S.R. and E.R. conducted the fieldwork. S.R. and F.T. analyzed the data. S.R. wrote the original draft, and the final version was produced jointly by all coauthors. We thank R. Stern and J. Muller for providing insights on developing these study methods. We thank E. Ramati for assisting with conducting the measurements by the mobile lab, and E. Schwartz for processing the long-term flux-tower and meteorological data. This project was supported by: ISF grants no. 2579/16, ISF grants no. 1976/17, And the JNF-KKL. Data is deposited in the European Fluxnet Database (http://europe-fluxdata.eu). Meteorological data are available from the Israeli Meteorological Service IMS (https://ims.data.gov.il/). Additional data is deposited in the lab archive and available from the corresponding authors.

