## Supplemental Table S1 and Figure S1 for "Large variations in afforestation-related climate cooling and warming effects across short distances"

**Table S1.** Study sites descriptions. Geographical and vegetation information for the three study sites across the Israeli climatic gradient used in this research.

| Site | Yatir | Eshtaol | Bir | Oak Forest |
| --- | --- | --- | --- | --- |
| Climatic classification | Arid (A) | Semi-Aridya (SA) | Dry Sub-Humid (DSH) | Semi-Arid (SA) |
| Latitude coordinate  Longitude coordinate | 31° 20' 49.20” N  35° 03' 07.20” E | 31° 47' 34.50” N  35° 00' 11.50” E | 33° 00' 00.50” N  35° 30' 40.50” E | 32° 44' 47.00" N  35° 13' 55.70" E |
| Mean annual rainfall (mm) | 285 | 543 | 755 | 580 |
| Daily mean temperature (°C) | 18 | 19 | 17 | 21 |
| Elevation (m) | 650 | 380 | 755 | 190 |
| Forest ecosystem  *(dominant species)* | Pine forest plantation  (*Pinus halepensis*) | Pine forest plantation  (*Pinus halepensis*) | Pine forest plantation  (*Pinus halepensis*) | Oak native forest  (*Quercus calliprinos* & *Q.ithaburensis (* |
| Trees age (y) | 50 | 50 | 40 | ~ 70 |
| Forest density (t ha^-1^) | 300 | 450 | 600 | 280 |
| Canopy height (m) | 10 | 11 | 11 | 8 |
| Soil type | Light brown Rendzina | Light brown Rendzina | Rendzina and Terra rossa | Rendzina and Terra rossa |
| Shrubland ecosystem | Grass under heavy grazing | Mixed shrubs  (annual vegetation) | Grass under heavy grazing | - |

**Figure S1.** **Long-term (1950 – 2021) annual rainfall amount and their trend over time in the three study regions:** Birya - DSH, Eshtaul - SA, and Yatir - A. While the DSH and the SA sites show no clear P trend over the 70 records years, the arid site trend, 2.2 mm per decade, is small (8% P reduction) and is highly sensitive to the initiate and the ending year's precipitation amounts.

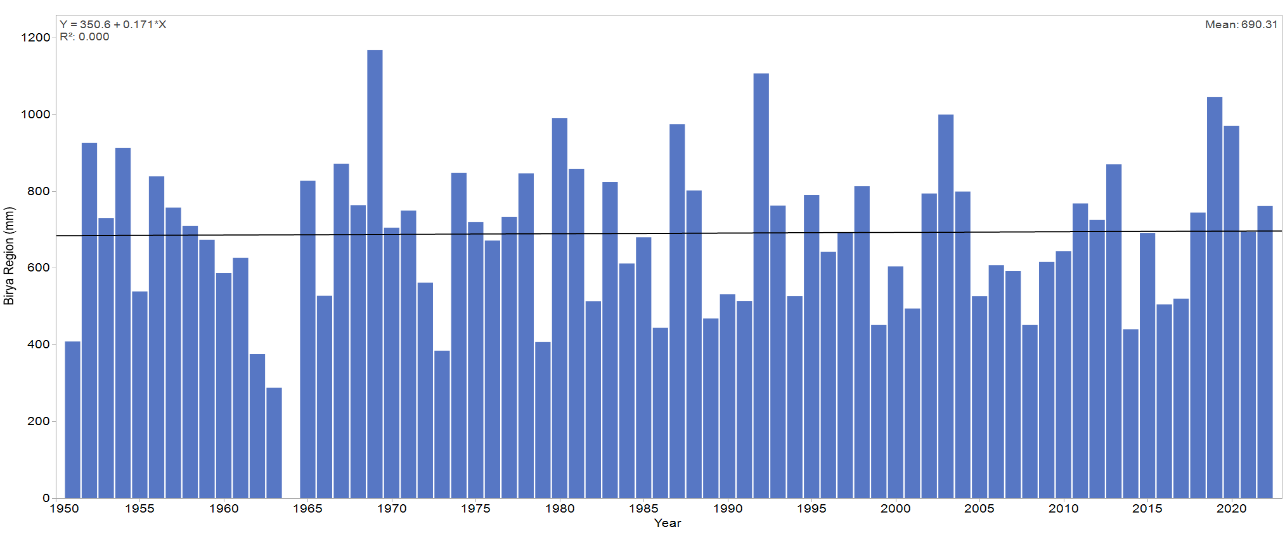

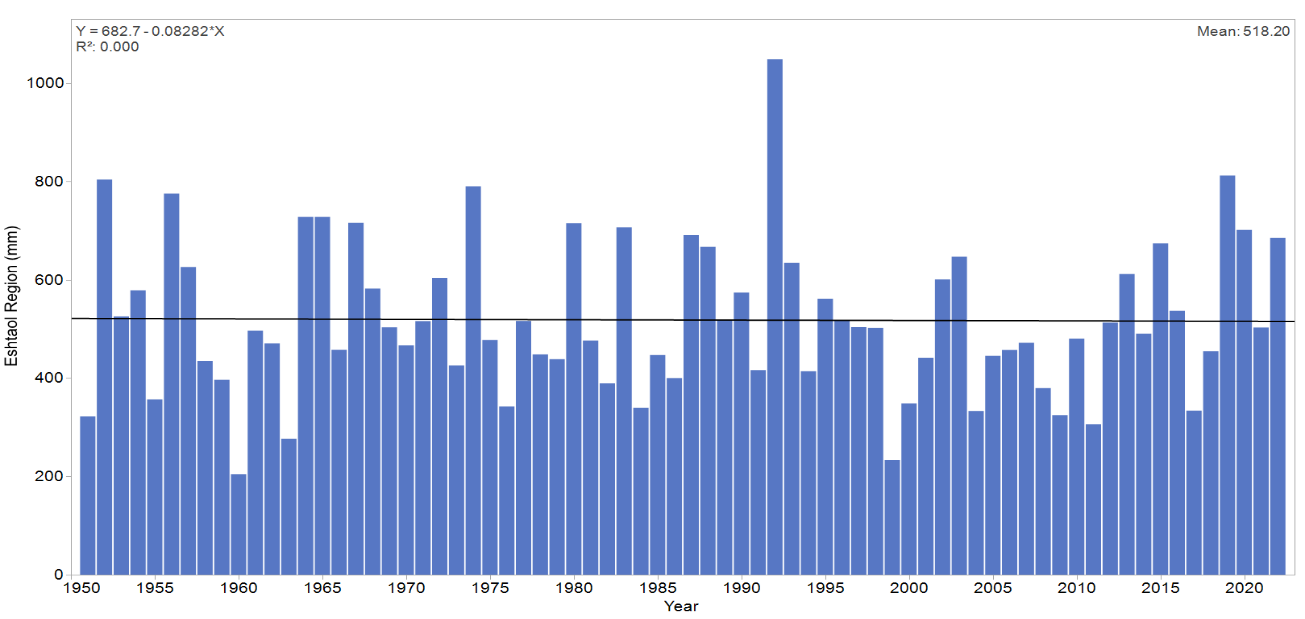

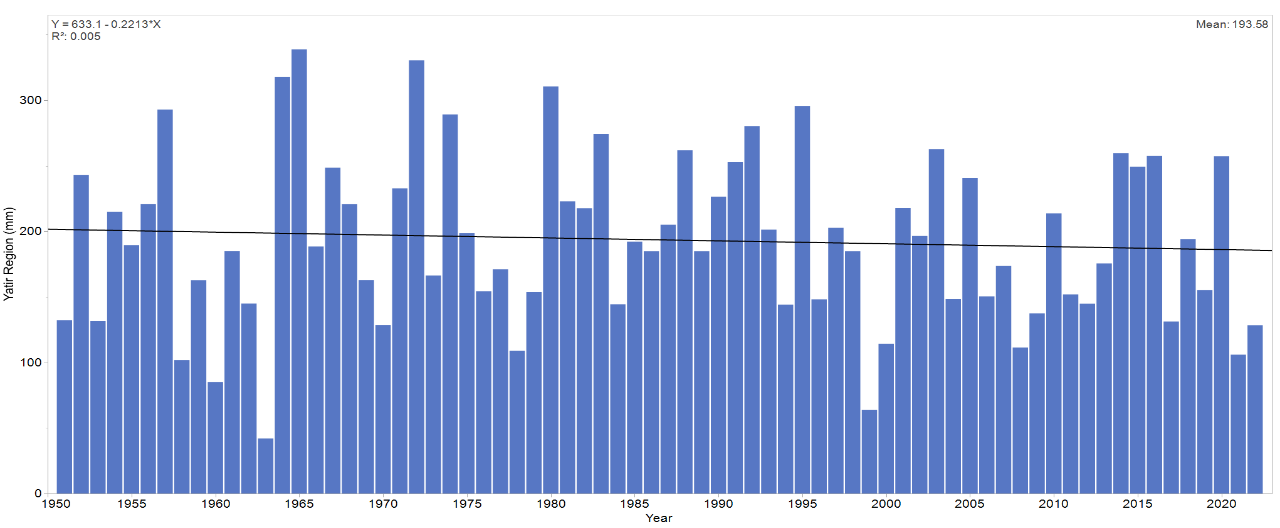

Annual P values were taken from close-by meteorological stations (Israel meteorological service, <https://ims.data.gov.il/he/ims/3>) having long-term records thus, study sites' P values could differ from the above P values.
